# Identification of an osteopontin-derived peptide that binds neuropilin-1 and activates vascular cells

**DOI:** 10.1101/2023.09.05.556237

**Authors:** Yihong Chen, Chrysostomi Gialeli, Junyan Shen, Pontus Dunér, Björn Walse, Annette Duelli, Rhawnie Caing-Carlsson, Anna M. Blom, John Zibert, Anna Hultgårdh Nilsson, Jan Alenfall, Chun Liang, Jan Nilsson

## Abstract

**Background:** The osteopontin-derived peptide FOL-005 has been shown to stimulate hair growth. Using ligand-receptor glycocapture technology we identified neuropilin-1 (NRP-1), a known co-receptor for vascular endothelial growth factor (VEGF) receptors, as the most probable receptor for FOL-005. Considering that induction of perifollicular angiogenesis by VEGF is a critical step in activation of the anagen growth phase of hair follicles, we investigated the effect of the more stable FOL-005 analogue FOL-026 on vascular cells.

**Methods:** X-ray diffraction and microscale thermophoresis analysis were used to determine receptor binding of FOL-peptides. Cultured human arterial smooth muscle cells were used for studies of cell proliferation, migration, intracellular signaling, and apoptosis. A Matrigel plug assay was used to study angiogenesis in vivo. RNA seq. was used to analyze changes in gene expression patterns.

**Results:** FOL-026 was found to share binding site with VEGF in the NRP-1 b1-subdomain. Stimulation of human umbilical vein endothelial cells with FOL-026 resulted in phosphorylation of AKT and ERK1/2, increased cell growth and migration, stimulation of endothelial tube formation and inhibition of apoptosis *in vitro,* as well as activation of angiogenesis *in vivo.* Down-regulation of NRP-1 by NRP-1-specific small interfering RNA blocked the stimulatory effects of FOL-026 on endothelial cells. RNA sequencing showed that FOL-026 activated pathways involved in tissue repair. Exposure of human coronary artery smooth muscle cells to FOL-026 stimulated cell proliferation, migration, inhibited apoptosis, and induced VEGF gene expression by an NRP-1-dependent mechanism.

**Conclusions:** These findings identify NRP-1 as the receptor for FOL-026 and show that its biological effects mimic that of growth factors binding to the VEGF receptor family. They also suggest that FOL-026 may have therapeutical potential in conditions that require vascular repair and / or enhanced angiogenesis.

**Clinical perspective:**

- Insufficient vascular repair increases the risk of acute cardiovascular events, something that is of particular importance in diabetes.
- A peptide derived from the extracellular matrix protein osteopontin (FOL-026) binds to the vascular endothelial growth factor (VEGF) co-receptor neuropilin-1 stimulating endothelial and vascular smooth muscle cell proliferation and migration as well as the formation of new vessels.
- By mimicking the effects of VEGF FOL-026 may have therapeutical potential in conditions that require vascular repair and / or enhanced angiogenesis such as vascular complications to diabetes, peripheral artery disease and salvage of ischemic myocardium.

## 1. Introduction

Osteopontin is an extracellular matrix protein consisting of 314 amino acids and with a molecular weight ranging from 41 to 75 kDa depending on different forms of post-translational modifications.^1^ It is widely expressed in different organs throughout the body playing important roles in the interaction between cells and the surrounding extracellular matrix.^2,3^ Osteopontin affects many physiological processes including bone mineralization, wound healing, and inflammation as well as pathological conditions such as insulin resistance and tumor development. The diversity of effects is likely explained by different post-translation modifications in different organs and pathological processes, but also by the presence of multiple binding sites including the C terminal binding to CD44, the N terminal integrin binding site and a cryptic RGD-containing integrin binding site that is exposed in response to thrombin cleavage.^2^ We have previously found that a peptide derived from the latter binding site, but with point mutations in the RGD sequence, stimulates hair growth in an animal model by promoting the transition of hair follicles from the telogen to the anagen phase.^4^ The effect of this peptide (FOL-005) on hair growth has been investigated in three clinical trials (NCT02793557, NCT03467412, and NCT04774874) providing support for the presence of stimulatory hair growth activity also in humans, and with a good safety profile.

Hair growth is regulated by the cycling of the hair follicle from telogen (resting) phase to the anagen (growth) phase and finally to the catagen (regression) phase.^5,6^ During the anagen phase there is an extensive proliferation of follicular keratinocytes along with an elongation and thickening of the hair shaft. Resting hair follicles are avascular but during the anagen phase there is an ingrowth of blood vessels from the deeper layers of the dermis to meet the nutritional needs of the active hair follicle.^7^ Alopecia is characterized by decreased perifollicular vascularization.^8^ Studies by Yano et al^9^ demonstrated that the increased vascularization of hair follicles during the early anagen phase was associated with an upregulation of vascular endothelial growth factor (VEGF) mRNA in the follicular keratinocytes of the outer root sheath. Transgenic overexpression of VEGF in these cells stimulated perifollicular vascularization and hair growth in mice. Moreover, systemic treatment with a VEGF neutralizing antibody resulted in reduced perifollicular vascularization and hair growth. Several other studies have subsequently confirmed the role of locally expressed VEGF, as well as hepatocyte growth factor (HGF), another important endothelial mitogen, in hair growth.^10–13^ Against this background, we set out to investigate the cellular binding site of FOL-005 and to determine its effects on cultured human vascular cells. The latter studies were performed using FOL-026 (VDTYDG**<u>G</u>**ISVVYGLR), a derivative of FOL-005 (VDTYDG**<u>D</u>**ISVVYGLR), in which one aspartic acid has been substituted with glycine to reduce the risk of deamidation and increase the stability of the peptide.

## 2. Materials and methods

### 2.1 ELISA binding assays

Maxisorp 96 well microtiter plates (Thermo Fisher Scientific, NY) were coated with 8μg/ml of human neuropilin-1 (NRP-1, Acro systems, Newark, DE) or 10μg/ml of human NRP-1 structural variants or heparin. Unspecific binding was blocked with 1% BSA. Biotin labeled FOL-026 was added to the plates overnight. The interaction was detected with streptavidin-HRP and OPD substrate.

### 2.2 Microscale thermophoresis

Human NRP-1 (Phe22-Lys6444, Acro systems, Newark, DE, USA), and purified NRP-1 b1, NRP-1 b1b2 domains were labeled using the Protein Labeling Kit RED-NHS (NanoTemper Technologies, Munich, Germany). The interaction of NRP-1 structural variants with FOL-peptides was measured using a Monolith NT.115 [NT.115Pico/NT.LabelFree] instrument (NanoTemper Technologies, Munich, Germany). For more details see supplementary materials and methods.

### 2.3 Crystallization and modelling

A detailed description of the experimental approach concerning the purification of NRP-1 structural variants, crystallization, freezing and data collection and refinement are provided in Supplementary Materials and Methods and Supplementary Table 1.

### 2.4 Preparation of peptides

The peptide FOL-005 (sequence: VDTYDGDISVVYGLR, Mw: 1672 Da) was produced (PolyPeptide Laboratories, Strasbourg, France and Torrance, CA). The FOL-026 peptide (sequence: VDTYDGGISVVYGLR, Mw: 1614 Da) and the biotinylated FOL-026 (sequence: Biotin-VDTYDGGISVVYGLR, Mw: 1840 Da) were purchased from Biopeptide (San Diego, CA). All peptides were delivered as >95% pure as controlled by HPLC. The modification of the osteopontin sequence consists of the removal of two amino acids and insertion of one amino acid in the integrin-binding motif Arg-Gly-Asp (RGD) site of osteopontin which alters the functional RGD binding site of osteopontin.

### 2.5 Cell culture

Human umbilical vein endothelial cells (HUVECs) and human coronary artery smooth muscle cells (HCASMCs) were purchased from Thermo Fisher Scientific (Waltham, MA) and QuiCell (Shanghai, China). HUVECs were grown in medium 200 (Gibco, Grand Island, NY) containing 2% low serum growth supplement (LSGS, Gibco, Grand Island, NY) or endothelial cell medium (ECM, ScienCell, Carlsbad, Ca) supported with 5% fetal bovine serum (FBS, Gibco, Thornton, Australia) as well as 1% endothelial cell growth supplement (ECGS, ScienCell, Carlsbad, Ca). HCASMCs were cultured in medium 231 (Gibco, Grand Island, NY) containing 5% smooth muscle growth supplement (SMGS, Gibco, Grand Island, NY) or smooth muscle cell medium (SMCM, ScienCell, Carlsbad, CA) supplemented with 2% FBS and 1% smooth muscle cell growth supplement (SMCGS, ScienCell, Carlsbad, CA). Antibiotic-Antimycotic supplement (1%, Gibco, Grand Island, NY) were always added to the culture medium. Lyophilized FOL-026 was dissolved in PBS (Gibco, Paisley, UK) and diluted to the indicated concentrations with corresponding culture medium.

### 2.6 Cell starvation

Cells were seeded with the density required for different experiments. Before peptide stimulations, HUVECs were incubated in the medium 200 plus 0.5% supplement or ECM mixed with 0.5%FBS and 0.1% ECGS for 24 hours. HCASMCs were starved in medium 231 or SMCM without any supplements for 24 hours. Cell starvation was not performed in all cell experiments with small interfering RNA.

### 2.7 Cell transfection

Small interfering RNA against NRP-1 (siNRP-1) and the negative control Non-targeting siRNA (siRNA-NC) were purchased from Dharmacon (Lafayette, CO). For cell transfection, siNRP-1/siRNA-NC, serum-free medium and HiPerFect Transfection Reagent (Qiagen, Hilden, Germany) were premixed and incubated with cells according to manufacturer’s instructions. All assays were carried out 48-72 hours after transfection.

### 2.8 Cell proliferation assay

Cell proliferation was analyzed with BrdU Cell Proliferation Elisa Kit (Abcam, Cambridge, UK). In brief, cells on 96-well plates treated with the FOL-026 for 32 hours, then 20μl mixture of BrdU reagent and culture medium was added into each well. After 16 hours of incubation, cells were fixed, and subsequent steps were performed according to the manufacturer’s instructions. The colorimetric reaction was read by Wallac 1420 Victor 2 (Perkin Elmer, Waltham, MA) or Synergy H1 microplate reader (BioTek, Winooski, VT) at 450nm.

### 2.9 Apoptosis and oxidative stress assay

Caspase-Glo 3/7 Assay (Promega, Fitchburg, WI) and ROS-Glo H_2_O_2_ Assay (Promega, Fitchburg, WI) were used to evaluate cell apoptosis and reactive oxygen species (ROS) generation. Initially, HUVECs and HCASMCs were seeded into 96-well plates at a density of 3-5 x 10^3^ per well and cultured overnight. After 48 hours of peptide pre-treatments, cells were exposed to a low-oxygen condition at 0.1% O_2_ (Smartor 118, Hariolab, Guangzhou, China) or 50μg/ml oxidized low-density lipoprotein (oxLDL, Invitrogen, Eugene, OR) for 24 hours and 2 hours, to detect active caspase-3/-7 and ROS signals. For analysis of apoptosis induced by soluble Fas ligand (sFasL, PeproTech, NJ,), cells were incubated with FOL-026 peptide for 24 hours before adding 5.0μg/ml sFasL and measurement of active caspase-3/-7 was performed at 48 hours. Subsequent steps were performed according to the manufacturer’s instructions. Luminescence signal was read by Wallac 1420 Victor 2 (Perkin Elmer, Waltham, MA) or Synergy h1 microplate reader (BioTek, Winooski, VT).

### 2.10 Tube formation assay

HUVECs on 6-well plates were starved for 24 hours or for siRNA transfection for 48 hours and dissociated by TrypLE Express (Gibco, Grand Island, NY) to obtain cell suspensions. Cells were seeded at a density of 1.5 x 10^4^ / well in 96-well plate pre-coated with Geltrex reduced growth factor basement membrane matrix (Gibco, Grand Island, NY) in Medium 200 complete culture medium containing different concentration of FOL-026 peptide. After 12-18 hours of incubation, random images were recorded by an inverted microscope (Nikon, Eclipse TE2000-U, Kawasaki, Japan). Total length, total master segments length, total branching length and total segments length were quantified using Angiogenesis Analyzer plug-in ImageJ software program (pixel/unit). Data were quantified with images from 7-12 wells per group.

### 2.11 In vivo Matrigel plug assay

All animal experiments were approved by the Committee on Ethics of Medicine at Naval Medical University and carried out in compliance with the guidelines of National Institutes of Health guide for the care and use of laboratory animals. Male BALB/c nude mice with an age of 8 weeks were purchased from GemPharmatech Co, Ltd (Nanjing, China). All animals were housed in pathogen-free conditions with 12 hours light/dark cycle and had free access to autoclaved chow and water. Briefly, mice were anesthetized with oxygen-carried isoflurane. A mixture containing 1 x 10^6^ HUVECs pre-treated with different concentration of FOL-026 peptide and 500ul growth factors-reduced Matrigel (ABW Matrigengel, Shanghai, China) was injected subcutaneously into the flank of mice, resulting in the formation of a solid plug *in vivo*. After 14 days, the mice were sacrificed with isoflurane overdose and the plugs were taken out for further analysis. For immunofluorescence staining of CD31(also known as platelet and endothelial cell adhesion molecule 1, PECAM-1), the samples were fixed in 4% formaldehyde solution overnight and embedded in paraffin. Sections were deparaffinized, rehydrated in xylene and a graded ethanol series, followed by a heat-based antigen retrieval. After blocking with goat serum, sections were incubated with CD31 primary antibody (Rb, Servicebio, Wuhan, China) at 4 °C overnight. Next day, a Cy3-conjugated goat anti-rabbit IgG antibody (Jackson ImmunoResearch, West Grove, PA) was used as secondary antibody for 1 hour at room temperature and nuclei were counterstained with DAPI (Invitrogen, Eugene, OR). For immunohistochemical staining of CD31, 3% H_2_O_2_ was used to suppress endogenous peroxidase activity before serum blocking. After incubation with primary and secondary (HRP-conjugated goat anti-rabbit IgG antibody, Servicebio, Wuhan, China) antibodies, DAB Substrate (Thermo Scientific, Rockford, IL) was used to achieve peroxidase reaction followed by hematoxylin (Sigma, St. Louis, MO,) counterstain for visualization. CD31 staining area was analyzed using image J software.

### 2.12 In vitro wound closure assay

Cells with exposure of starvation for 24 hours or siRNA transfection for 48 hours were seeded at a density of 3.5 x 10^4^/ inlet into Culture-Insert 2 well (Ibidi, Fitchburg, WI) in 24-well plate and cultured overnight. In the second day, the cell-free gap between two inlets was created by removing the insert and cells were washed by DPBS gently to remove the debris. Culture medium containing 2 % supplements and different concentration of FOL-026 peptide was added and images recorded with an inverted microscope (Nikon Eclipse TE2000-U, Kawasaki, Japan or Olympus CKX53, Tokyo, Japan) at different time-points. The percentage of wound closure was calculated using MRI-Wound-Healing plug-in ImageJ software program: wound closure % =1 − (wound area at *T*_5_ / wound area at *T*_0_)] × 100 (*T*_0_ is the wound area at 0 h, and *T*_5_ is the wound area at 5 hours after creating the gap (pixel/unit). Data were quantified with images from 8-10 wells per group.

### 2.13 Western blot

Cell samples were washed using pre-cold PBS and lysed with RIPA lysis buffer including protease and phosphatase inhibitor cocktail (Beyotime Biotechnology, Shanghai, China) on ice. The protein content of sample homogenate was quantified by the BCA Protein Assay Kit (Beyotime Biotechnology, Shanghai, China). After that, the mixture of sample and loading buffer was boiled for 10mins to denature. A 20ug aliquot of protein sample from each group was loaded onto SDS-PAGE gels and separated via electrophoresis, after which it was transferred to PVDF membranes (Millipore, Burlington, MA). Membranes were blocked with 5% BSA in TBS containing 0.1% Tween 20 (TBST) and incubated subsequently with primary antibodies including phospho-AKT (Thr308 from Signalway Antibody, College Park, MD and Ser473 from Proteintech, Rosemont, IL), AKT (Proteintech, Rosemont, IL), p42/44 MAPK (ERK1/2, Cell Signaling Technology, Danvers, MA), phosphor-p42/44 MAPK (p-ERK1/2, Cell Signaling Technology, Danvers, MA) and GAPDH (Cell Signaling Technology, Danvers, MA) at 4°C overnight. Corresponding secondary antibodies (Abcam, Cambridge, UK) were used to incubate membranes at room temperature for 1 hour the following day. After washing three time, the signal of protein was enhanced by enhanced chemiluminescence (WesternBright ECL, Advansta, San Jose, CA) substrate and detected with Quickgel 6200 (Monad, Wuhan, China). Bands were quantified using image J software.

### 2.14 RNA isolation and quantitative real-time PCR (qRT-PCR)

Vascular cells in 6-well plates were washed with cold PBS after discarding the supernatant medium. Total RNA was extracted with Trizol reagent (Invitrogen, Carlsbad, CA) and PureLink RNA Mini Kit (Ambion, Grand Island, NY) according to manufacturer’s instructions. The nucleic acid quantification was determined using NanoDrop 2000c (Thermo Fisher Scientific, Waltham, MA). Reverse transcription and real-time qPCR reaction were carried out with KAPA SYBR FAST One-Step qRT-PCR Master Mix Kit (KAPA Biosystems, Wilmington, MA) on LighyCycler 480 system (Roche, Mannheim, Germany). All primers were purchased from miScript Primer Assays (Qiagen, Valencia, CA) and GAPDH mRNA expression of each sample was used as normalization.

### 2.15 RNA sequencing (RNA-seq) analysis

For RNA-seq, the protocol of total RNA extraction from vascular cells was done as described above. The RNA library was generated and sequenced with Illumina 2000 platform (LC Sciences, Houston, TX). Differentially expressed genes (DEGs) were screened with a threshold of |logFC|>0.5 and *p* value < 0.05. The Mfuzz package in R was used to group DEGs into 12 clusters with specific gene expression patterns. For Gene Ontology (GO) and Kyoto Encyclopedia of Genes and Genomes (KEGG) analysis, clusterProfiler package was used to achieve functional gene annotations and obtain the potential pathways regulated by peptide. Terms of GO enrichment analysis in biological processes and KEGG analysis were ranked with *p* value and the top 10 terms shown. For protein-protein interaction (PPI) analysis, Search Tool for the Retrieval of Interacting Genes (STRING) database was used to construct a PPI network and Cytoscape software with cytohubba plug-in was used to identify hub genes.

### 2.16 Statistical analyses

Statistical analyses were carried out with Prism 8 or 9 software. Data was presented as means ± standard deviation (SD) or as means ± standard error of mean (SEM) as specified. Experiments were analyzed using Student’s t test (between two groups) and one-way ANOVA with Dunnett’s multiple comparisons tests or two-way ANOVA with Sidak’s multiple comparisons tests (between multiple groups). A *p* value of <0.05 was considered significant and the respective p values in figures are indicated as follows: *P<0.05, **P<0.01, ***P<0.001, ****P<0.0001.

## 3. Results

### 3.1 FOL-005 and FOL-026 bind to NRP-1

Using ligand-receptor glycocapture technology we identified NRP-1 as the most probable receptor for FOL-005 on cultured human hair outer root sheet cells (Supplementary Fig. S1A). Using plates coated with purified NRP-1 we confirmed that FOL-026 also bound to NRP-1 and that this binding could be blocked by both VEGF and heparin (Fig. 1A). FOL-026 was also found to bind to heparin (Fig. 1B). The binding affinity between NRP-1 and fluorescently N-terminal labelled FOL-peptides was measured by detecting variations in fluorescence caused by an IR-laser induced temperature change, using microscale thermophoresis (MST) (Fig. 1C). The b1 and b2 domains of the of human NRP-1b1b2 in complex with FOL-026 show the typical jelly-roll β-barrel fold, with the peptide binding with its C-terminal arginine in the VEGF binding site (Fig. 1D). The most common interactions of ligands binding to the NRP-1 VEGF binding site are hydrogen bonds between D320 and a C-terminal arginine and hydrogen bonds between S346, Y353 and T349 to the backbone carbonyl group of the C-terminal arginine, as depicted in Figure 1E. The superposition of FOL-026 with various ligands in the VEGF binding site of NRP1 shows that FOL-026 binds in a similar way to NRP-1 as previously investigated ligands (VEGFA, VEGFB186), as well as a commercial inhibitor of NRP-1 (EG01377) (Fig. 1F). FOL-026 exhibited stronger binding affinity to the NRP-1b1b2 than to the -1b1 structural variant (Fig. 1G-I). Comparing the binding affinities of FOL-005 and FOL-026 showed that the latter had a higher binding affinity to NRP-1b1b2 (Supplementary Fig. S2A). At the same time no major differences in the binding affinity between the peptides were observed for NRP-1b1 (Supplementary Fig. S2B).

**Figure 1.**
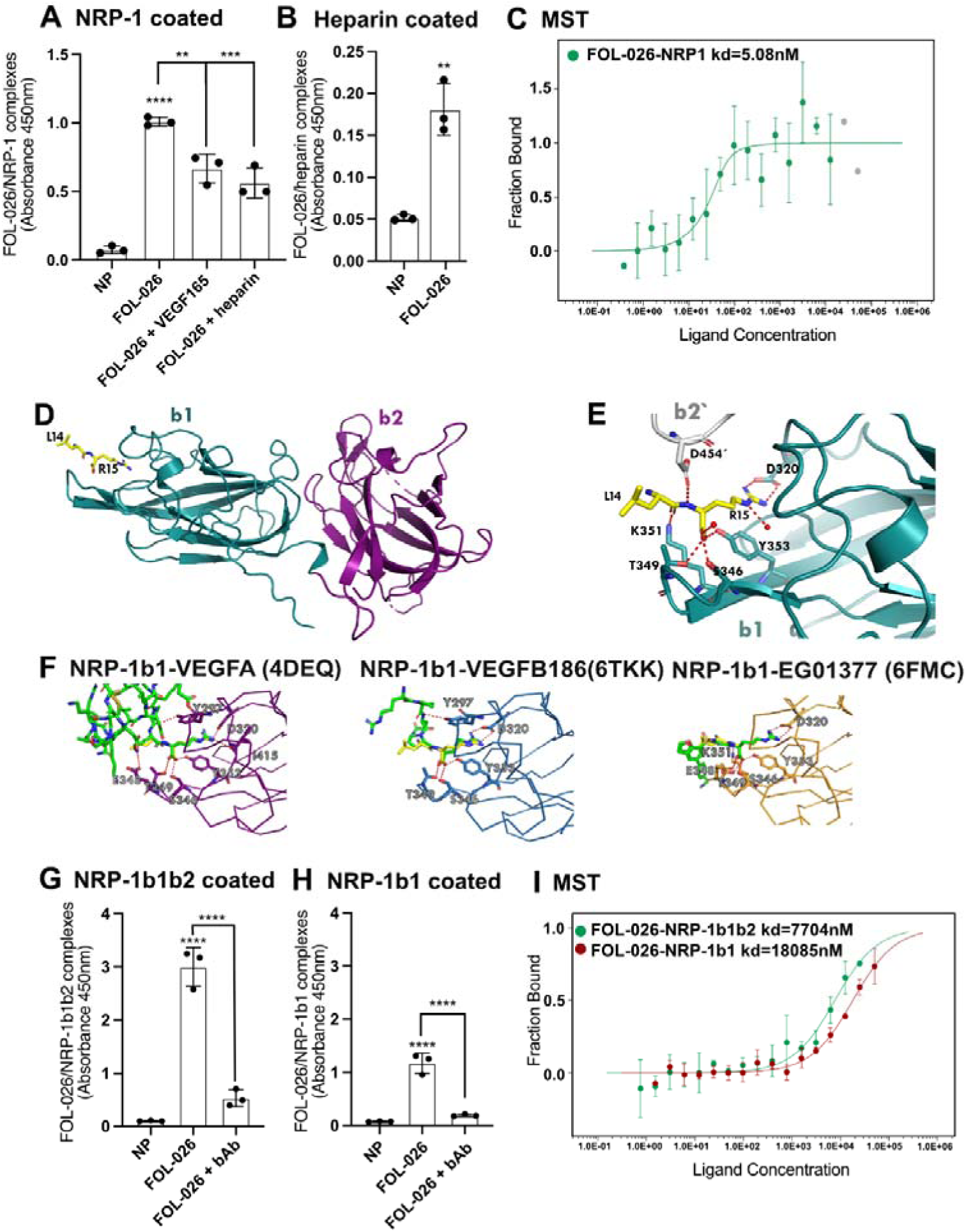
**(A)** Human NRP-1 coated plates (8μg/mL) were incubated overnight with 50μM N-terminal biotin labeled FOL-026 alone, or together with VEGFA165 (4.5μg/mL), or heparin (91μg/mL) and complexes of NRP-1/FOL-26 were monitored by a sandwich ELISA with Streptavidin-biotin detection. ND, no protein. **(B)** Human heparin coated plates (100μg/mL) were incubated overnight with 50μM biotin labeled FOL-026 and complexes of heparin/FOL-26 were monitored by a sandwich ELISA with Streptavidin-biotin detection. ND, no protein. **(C)** In this MST experiment the concentration of the labeled human NRP-1 was kept constant (50 nM), while the concentration of the non-labeled FOL-026 varied between 12.5 μM – 0.38 nM. After 20 min incubation at 37°C the samples were loaded into Monolith NT.115 [Premium] Capillaries (NanoTemper Technologies) and the MST measurement was performed using the Monolith NT.115 [NT.115Pico/NT.LabelFree] (NanoTemper Technologies) at 80% LED power and high MST power. MST traces with cold region start/end 0.25s/1.25s, hot region start/end 29.14s/30.14s was used for analysis, and a Kd of 5.08nM with Kd confidence +/-7.7 nM was derived for this interaction (n = 3 independent measurements, error bars represent the standard deviation). **(D)** Overview of the structure of NRP1 b1b2 (in green and purple) with FOL peptide (show in yellow sticks), **(E)** Close-up view of the bound peptide. Six different bonds are seen to D320, Se346, T349, Y353 and K351. One additional hydrogen bond is formed to the symmetry related D454’ and two hydrogen bonds to water, **(F)** Close-up view of the VEGF binding site with different ligands 4DEQ (purple), 6TKK (blue), 6FMC (orange) and structure with FOL026 peptide (green). **(G and H)** Human NRP-1 b1b2 coated plates were incubated overnight with 50μM biotin labeled FOL-026 alone, or together with 1μg/mL blocking NRP-1 Ab (bAb) and complexes of NRP-1/FOL-26 were monitored by a sandwich ELISA with Streptavidin-biotin detection. NP, no protein. **(I)** In this MST experiment the concentration of the labeled human NRP-1 structural variants were kept constant (20 nM), while the concentration of the non-labeled FOL-026 was varied between 50 μM – 1.53 nM. After 20 min incubation at 37°C the samples were loaded into Monolith NT.115 [Premium] Capillaries (NanoTemper Technologies) and the MST measurement was performed using the Monolith NT.115 [NT.115Pico/NT.LabelFree] (NanoTemper Technologies) at 40% LED power and high MST power. MST traces with cold region start/end -1s/0s, hot region start/end 9s/10s was used for analysis, and a FOL-026/NRP-1 b1b2 Kd of 7704.02 nM with Kd confidence +/-2763.5 nM, as well as a FOL-026/NRP-1 b1 Kd of 1885 nM with Kd confidence +/-3832.2 nM were derived for these interactions (n = 3 independent measurements, error bars represent the standard deviation). Unpaired t-test was used when comparing 2 samples, one-way ANOVA Dunnett’s multiple comparisons test was used when compared 3 or more samples. **P <0.01, ***P <0.001, and ****P <0.0001.

### 3.2 FOL-026 stimulates endothelial cell repair responses

To determine the effect of FOL-026 on endothelial cells, we used human umbilical vein endothelial cells (HUVECs). Exposure of HUVECs to 10nM and 100nM FOL-026 resulted in enhanced proliferation, although this effect was attenuated at a higher concentration (Fig.2A). Treatments with FOL-026 also rescued endothelial cells from apoptosis induced by low oxygen (Fig.2B) and sFasL (Supplementary Fig. S3A) with maximal effect at a concentration of 100nM. The luminescent signal of H_2_O_2_ from HUVECs exposed to oxLDL for 2 hours was analyzed for the assessment of ROS generation. Compared to control cells, pre-incubation with 10nM and 100nM FOL-026 enhanced the antioxidant capacity of HUVECs (Fig.2C). To investigate the impact of FOL-026 on migration we determined endothelial migration in an *in vitro* wound model (Fig.2D). FOL-026 stimulated HUVECs migration into the wound area by about 80% (Fig.2E) Treatment with 10nM of FOL-026 had no effect on VEGF-A gene expression (Fig.2F), while it reduced the VEGFR-2 (Fig.2G) and NRP-1 (Fig.2H) mRNA levels. At a concentration of 1000nM FOL-026 reduced VEGFR-2 mRNA expression (Fig.2G). Moreover, there was a down-regulation of tumor necrosis factor α (TNFα) mRNA expression (Supplementary Fig.S3B) in response to FOL-026, while there was no effect on interleukin 6 (IL-6) and matrix metalloproteinase-2 (MMP2) mRNA expression (Supplementary Fig.S3C and 3D). Interestingly, a concentration-dependent up-regulation of matrix metalloproteinase-3 (MMP3) expression was observed (Supplementary Fig.S3E). An association between MMP3 and angiogenesis was reported in a GWAS study.^14^

**Figure 2.**
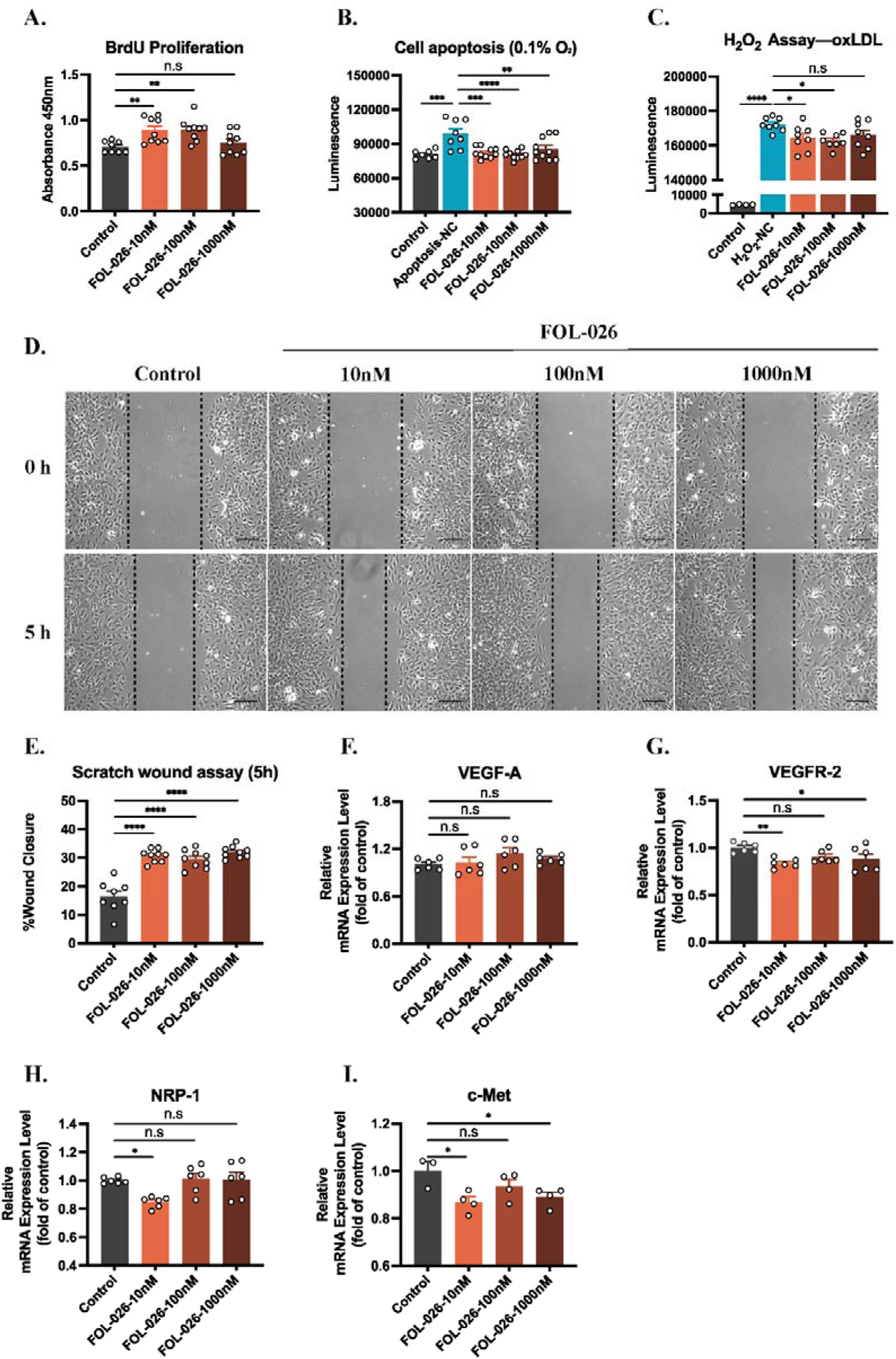
Effect of FOL-026 on endothelial cells. The impact of FOL-026 on human umbilical vascular endothelial cells (HUVECs) proliferation **(A)** and apoptosis **(B)** induced with low-oxygen condition was determined by BrdU uptake (n=8-9 per group) and active caspase-3/-7 (n=7-10 per group) respectively. The reactive oxygen species (ROS) level **(C)** activated by 50ug/ml oxidized low-density lipoprotein (oxLDL) for 2 hours in cells pre-incubated with FOL-026 was assessed by H_2_O_2_ measurement (n=4-8 per group). Representative pictures of endothelial migration into an *in vitro* injury model are shown **(D)** and the wound closure rate **(E)** in HUVECs stimulated with increasing concentration of FOL-026 was quantified by image J software (n=8-9 per group). The effect of FOL-026 on gene expression of vascular endothelial growth factor-A (VEGF-A, **F**) and its receptors **(G and H)**, as well as the hepatocyte growth factor (HGF) receptor c-Met **(I)** in HUVECs (n=3-6 for F-I). Scale bar=150μm in D. Data are presented as means±SEM and acquired from 3 to 4 independent replicate tests. *P<0.05, **P<0.01, ***P<0.001, and ****P<0.0001 by one-way ANOVA and Dunnett’s post-hoc test.

### 3.3 FOL-026 promotes angiogenesis and activates phosphorylation of PI3K/Akt and ERK1/2

To assess the effect of FOL-026 on angiogenesis, an endothelial tube formation assay was used (Fig.3A). The results showed that FOL-026 induced a significant increase in total length (Fig.3B), total master segments length (Fig.3C), total branching length (Fig.3D) and total segments length (Fig.3E) of endothelial tubes. Next, we used a subcutaneous Matrigel plug assay to investigate the proangiogenic effect of FOL-026 on human endothelial cells *in vivo* (Fig.3F). In line with the data from the tube formation assay, FOL-026 peptide led to a remarkable increase of neovascularization in Matrigel plugs (Fig.3G), which was confirmed by CD31^+^ immunofluorescence and immunohistochemical staining (Fig.3H and Supplementary Fig.S4A). Previous studies have reported that the Akt and ERK1/2 activation play vital roles in endothelial proliferation, wound healing and vessel formation.^15,16^ Consistent with this, western blot results showed that FOL-026 enhanced Akt and ERK1/2 phosphorylation in HUVECs in a concentration-dependent manner (Fig.3I-K). These observations show that FOL-026 promotes endothelial repair processes, angiogenesis, and actives the PI3K/Akt and ERK1/2 phosphorylation.

**Figure 3.**
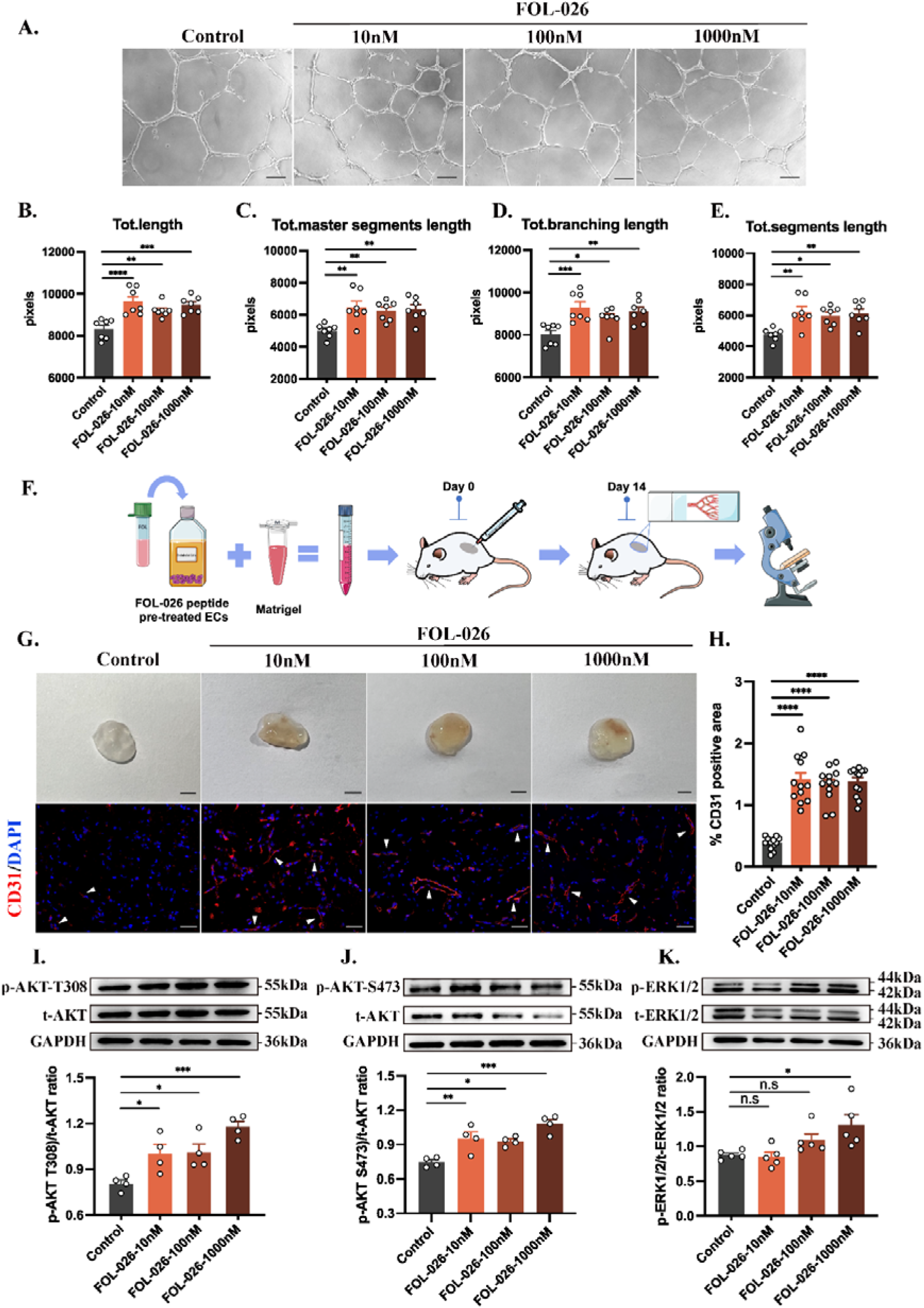
Effect of FOL-026 on angiogenesis in vitro and vivo. Representative images of tube formation in HUVECs treated with FOL-026 are shown **(A)**. Quantified measurement of total length **(B)**, total master segments length **(C)**, total branching length **(D)** and total segments length **(E)** were performed to evaluate angiogenesis *in vitro* (n=7 per group for B-E). The experimental flow of Matrigel plug assay *in vivo* is shown in **(F)**. Matrigel mixed with equal number of HUVECs pre-treated with indicated concentration of FOL-026 peptide were injected into the flank area of mice. Representative images of extracted Matrigel plugs (row 1) and CD31^+^ immunofluorescent (row 2) staining in Matrigel plug tissues **(G)**. Arrows indicate endothelial staining. The quantification of vessel density was assessed asCD31^+^ positive area in sections (**H**, n=12 per group). Scale bar=150μm in A, scale bar=50μm (row 1) and 200μm (row 2) in G. Western blot analysis of phosphorylated AKT (p-AKT-T308 and p-AKT-S473), t-AKT, phosphorylated ERK1/2 (p-ERK1/2) and t-ERK1/2 in endothelial cells stimulated with FOL-26 peptides for 48 hours **(I-K)**. Quantification of phosphorylation intensities normalized to the respective total protein (n=4 for I and J, n=5 for K). Data are presented as means±SEM and acquired from 3 to 4 independent replicate tests. *P<0.05, **P<0.01, ***P<0.001, and ****P<0.0001 by one-way ANOVA and Dunnett’s post-hoc test.

### 3.4 NRP-1 is a key mediator of the effects of FOL-026 on endothelial cells

Having established that FOL-026 binds to NRP-1, we next investigated the functional importance of NRP-1 binding by treating HUVECs with small interfering RNA (siRNA). Transfection with 1nM and 5nM NRP-1 siRNA (siNRP-1) in HUVECs for 48 hours reduced the NRP-1 mRNA expression by around 45% and 70%, respectively (Fig.4A). Accordingly, we used 5nM siRNA in further studies. Treatment with siNRP-1 and siRNA-NC did not affect cell proliferation and low oxygen-induced cell apoptosis in absence of FOL-026 (Fig.4B and 4C). However, the stimulation of endothelial cell proliferation and the enhanced resistance against apoptosis mediated by FOL-026 were inhibited by NRP-1 mRNA silencing (Fig.4B and 4C). Furthermore, suppression of NRP-1 mRNA expression not only reduced FOL-026-induced HUVECs migration in the *in vitro* wound model (Fig.4D and 4E), but also inhibited the endothelial tube formation induced by FOL-026 (Fig.5A) including the effects on total length (Fig.5B), total master segments length (Fig.5C), total branching length (Fig.5D) as well as total segments length (Fig.5E). These findings demonstrate that NRP-1 is the critical mediator of the effects of FOL-026 on endothelial cells.

**Figure 4.**
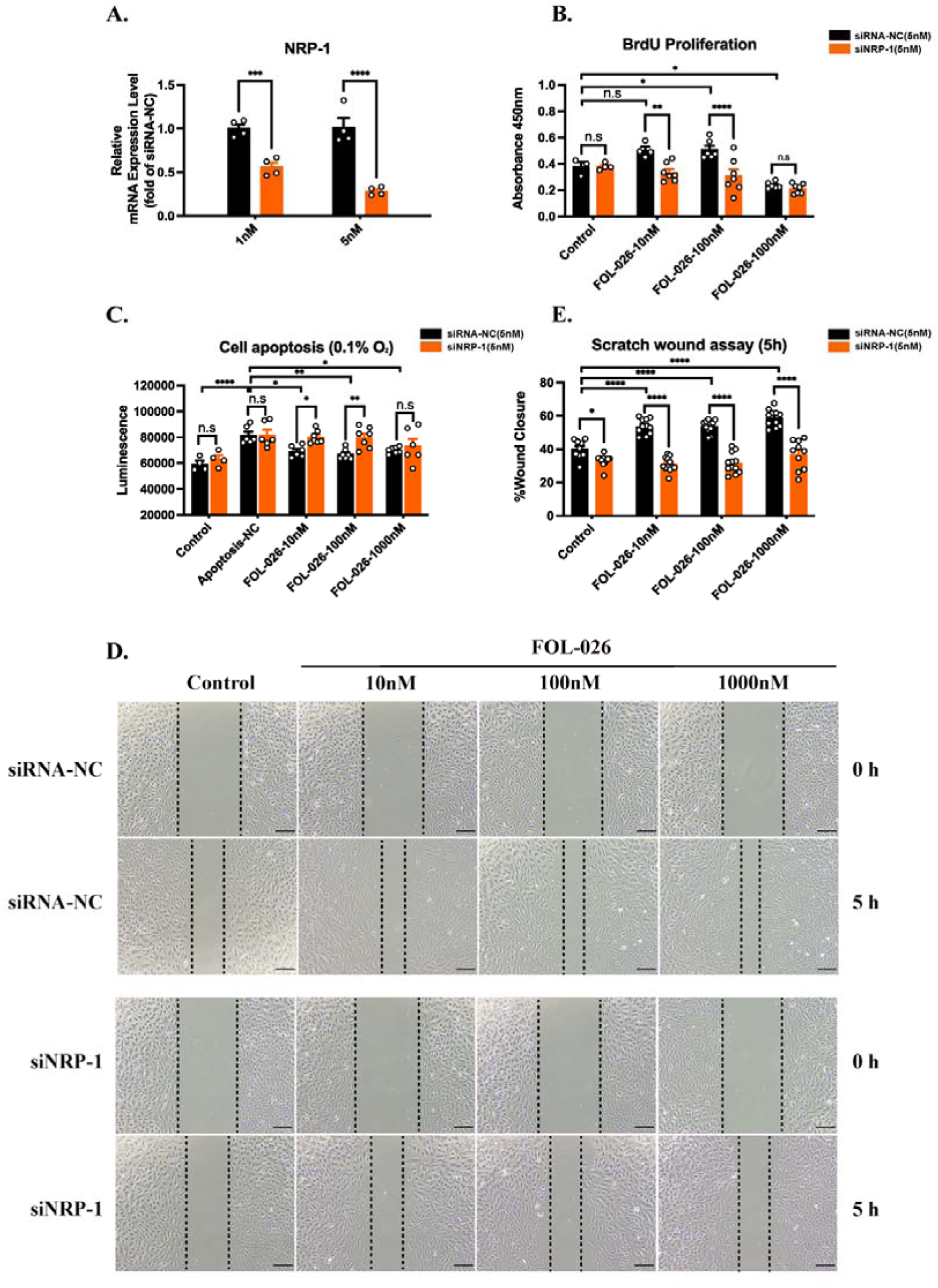
Effect of NRP-1 knock-down on endothelial cells stimulated with FOL-026. **(A)** The efficiency of small interfering RNA transfection for 48 hours in HUVECs was confirmed by qRT-PCR (n=4). The effect of FOL-026 stimulation on cell proliferation **(B)** and cell apoptosis at 0.1% oxygen **(C)** in HUVECs transfected with siRNA-NC or siNRP-1 was evaluated by BrdU incorporation (n=4-7 per group) and active caspase-3/-7 (n=4-7 per group). Representative pictures of endothelial cells at 0 and 5 hours after induction of the wound are shown **(D)**. The wound closure rate **(E)** of siRNA-NC/ siNRP-1-transfected HUVECs stimulated with increasing concentration of FOL-026 was quantified by image J software (n=9-12 per group). Scale bar =150μm in D. Data are presented as means ± SEM and acquired from 3 to 4 independent replicate tests. Two-way ANOVA followed with Sidak’s multiple comparisons test for A, two-way ANOVA followed with Sidak’s and Dunnett’s multiple comparisons test for B, C and E.*P<0.05, **P<0.01, ***P<0.001, and ****P<0.0001.

**Figure 5.**
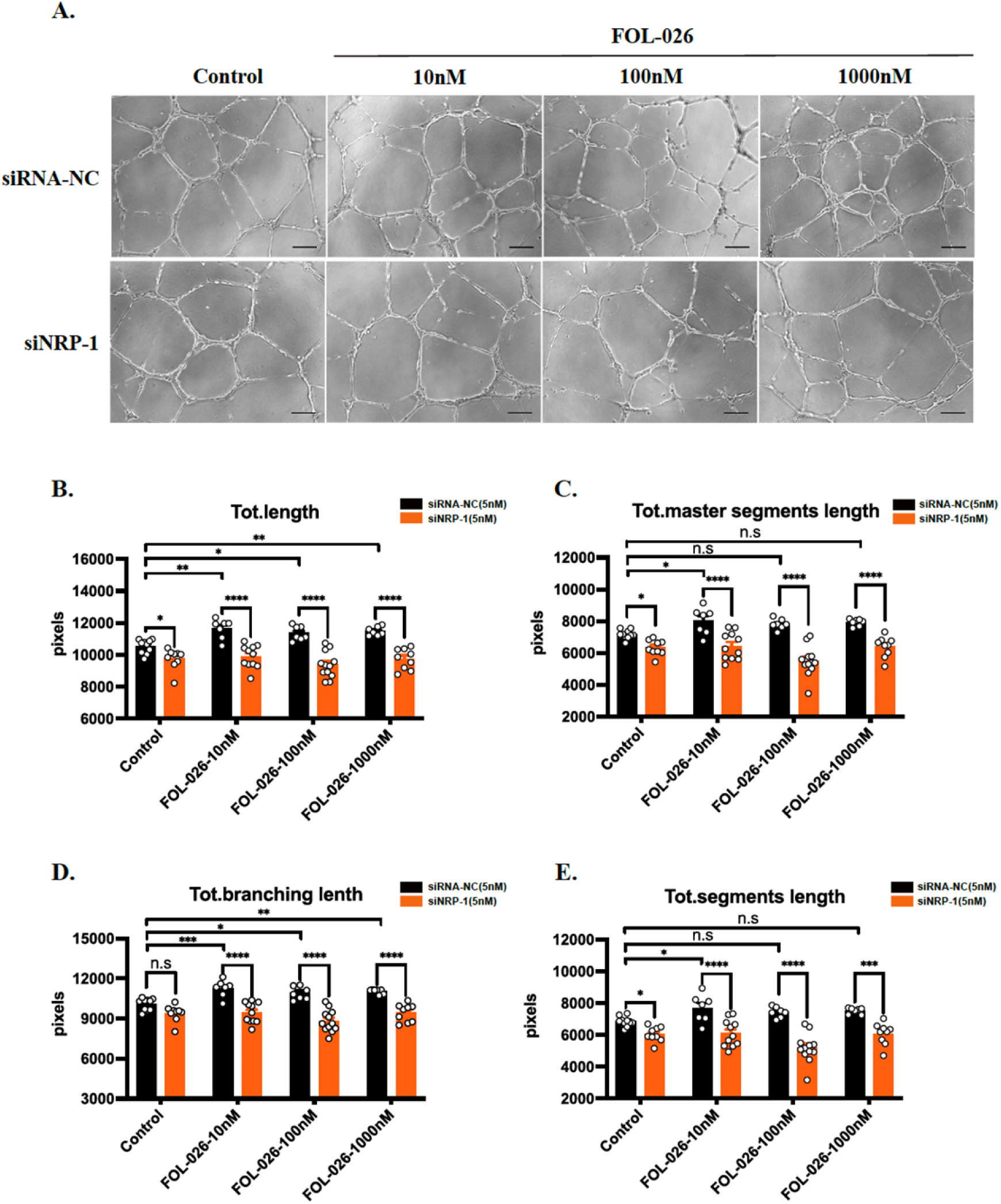
Effect of NRP-1 knock-down on tube formation induced by FOL-026. Tube formation with increasing concentrations of FOL-026 in the siRNA-NC and siNRP-1 groups are illustrated **(A)**, and total length **(B)**, total master segments length **(C)**, total branching length **(D)** as well as total segments length **(E)** were calculated using Angiogenesis Analyzer plug-in ImageJ (n=7-12 per group). Scale bar =150μm in A. Data are presented as means ± SEM and acquired from 3 to 4 independent replicate tests. *P<0.05, **P<0.01, ***P<0.001, and ****P<0.0001 by two-way ANOVA followed with Sidak’s and Dunnett’s multiple comparisons test.

### 3.5 FOL-026 stimulates arterial smooth muscle cell repair responses

To study the effect of FOL-026 on vascular smooth muscle cells, we used HCASMCs. Incubation with FOL-026 promoted HCASMCs proliferation with an attenuated effect at higher concentrations (Fig.6A). In line with the data from HUVECs, exposure of HCASMCs to 0.1% oxygen condition and 5 μg/ml sFasL activated apoptosis, and this was inhibited by pretreatment with FOL-026 (Fig.6B and Supplementary Fig. S5A). Moreover, FOL-026 pretreatment reduced the oxidative stress induced by exposure to oxLDL (Fig.6C). Stimulating HCASMCs with FOL-026 enhanced the *in vitro* wound closure rate at 10nM and 100nM (Fig.6D and 6E) and enhanced the mRNA expression of collagen type I α1 (COL1A1, Fig.6F) and α2 (COL1A2, Fig.6G). In addition, FOL-026 increased the gene expression of VEGF-A (Supplementary Fig. S5B), while reducing the mRNA level of VEGFR-2 and NRP-1 (Supplementary Fig. S5C and 5D). FOL-026 had no effect on gene expressions of the HGF receptor c-Met, TNFα, IL-6, and MMP2 (Supplementary Fig.S5E-5H), but upregulated gene expression of MMP3 at higher concentrations (Supplementary Fig.S5I). Next, we explored whether PI3K/Akt and ERK1/2 pathway were induced in smooth muscle cells stimulated with FOL-026. Consistent with the observations in endothelial cells, FOL-026 treatment caused AKT phosphorylation in smooth muscle cells, however, the effect was attenuated at the highest concentration of FOL-026 (Fig.6H and 6I). In contrast to the activation of AKT pathway, FOL-026 did not stimulate ERK1/2 phosphorylation in HCASMCs (Fig.6J). Collectively, these findings indicate that FOL-026 enhances the arterial smooth muscle cell repair responses, suppresses cell apoptosis, and stimulates PI3K/Akt phosphorylation in HCASMCs.

**Figure 6.**
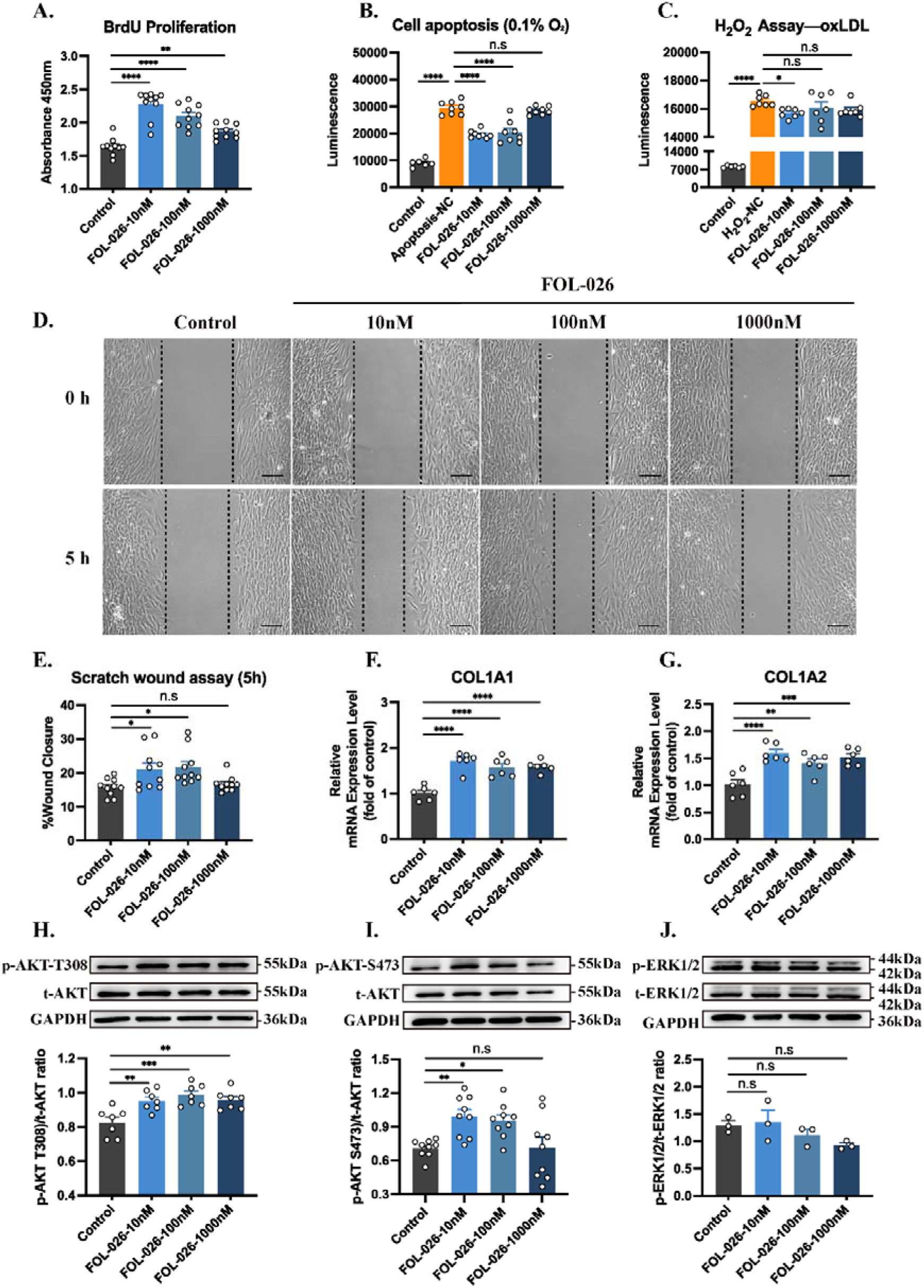
Effect of FOL-026 on arterial smooth muscle cells. The effect of FOL-026 treatment on smooth muscle cell proliferation **(A)** and apoptosis **(B)** induced by low-oxygen condition were evaluated with the uptake of BrdU (n=10 per group) and active caspase-3/-7 (n=6-8 per group) respectively. H_2_O_2_ measurement **(C)** in smooth muscle cells pre-incubated with FOL-26 peptides following 50μg/ml oxLDL exposure for 2 hours (n=6-7 per group). Representative images of the *in vitro* wound assay are shown **(D)** and the ability of wound closure **(E)** in human coronary smooth muscle cells (HCASMCs) treated with different concentrations of FOL-026 was measured by image J software (n=10 per group). qRT-PCR analysis of collagen type I α 1 chain (COL1A1, **F**) and collagen type I α 2 chain (COL1A2, **G**) mRNA expression in FOL-026 peptide-stimulated HCASMCs (n=6 for F and G). Western blot analysis of phosphorylated AKT (p-AKT-T308 and p-AKT-S473), t-AKT, phosphorylated ERK1/2 (p-ERK1/2) and t-ERK1/2 in HCASMCs stimulated with FOL-026 **(H-J)**. Quantification of Akt and ERK1/2 phosphorylation were normalized to respective total protein expressions (n=7 for H, n=9 for I, n=3 for J). Scale bar =150μm in D. Data are presented as means ± SEM and acquired from 3 to 4 independent replicate tests. *P<0.05, **P<0.01, ***P<0.001, and ****P<0.0001 by one-way ANOVA and Dunnett’s post-hoc test.

### 3.6 NRP-1 is a key regulator of FOL-026 effects on smooth muscle cells

To determine if NRP-1 binding contributed to the effects of FOL-026 on HCASMCs we suppressed NRP-1 expression by pre-incubation with 10nM NRP-1-specific siRNA (Fig.7A). This was found to completely inhibit the stimulatory effects of FOL-026 on HCASMCs proliferation (Fig.7B) as well as the protective effect against apoptosis (Fig.7C). Furthermore, suppression of NRP-1 mRNA expression inhibited FOL-026-stimulated HCASMCs migration in the *in vitro* wound model (Fig.7D-7E). Interestingly, the silencing of NRP-1 was observed to reduce HCASMCs migration also in the absence of FOL-026. This is most likely explained by a role of NRP-1 in the autocrine loop of placental growth factor-stimulated HCASMCs migration previously described.^17^

**Figure 7.**
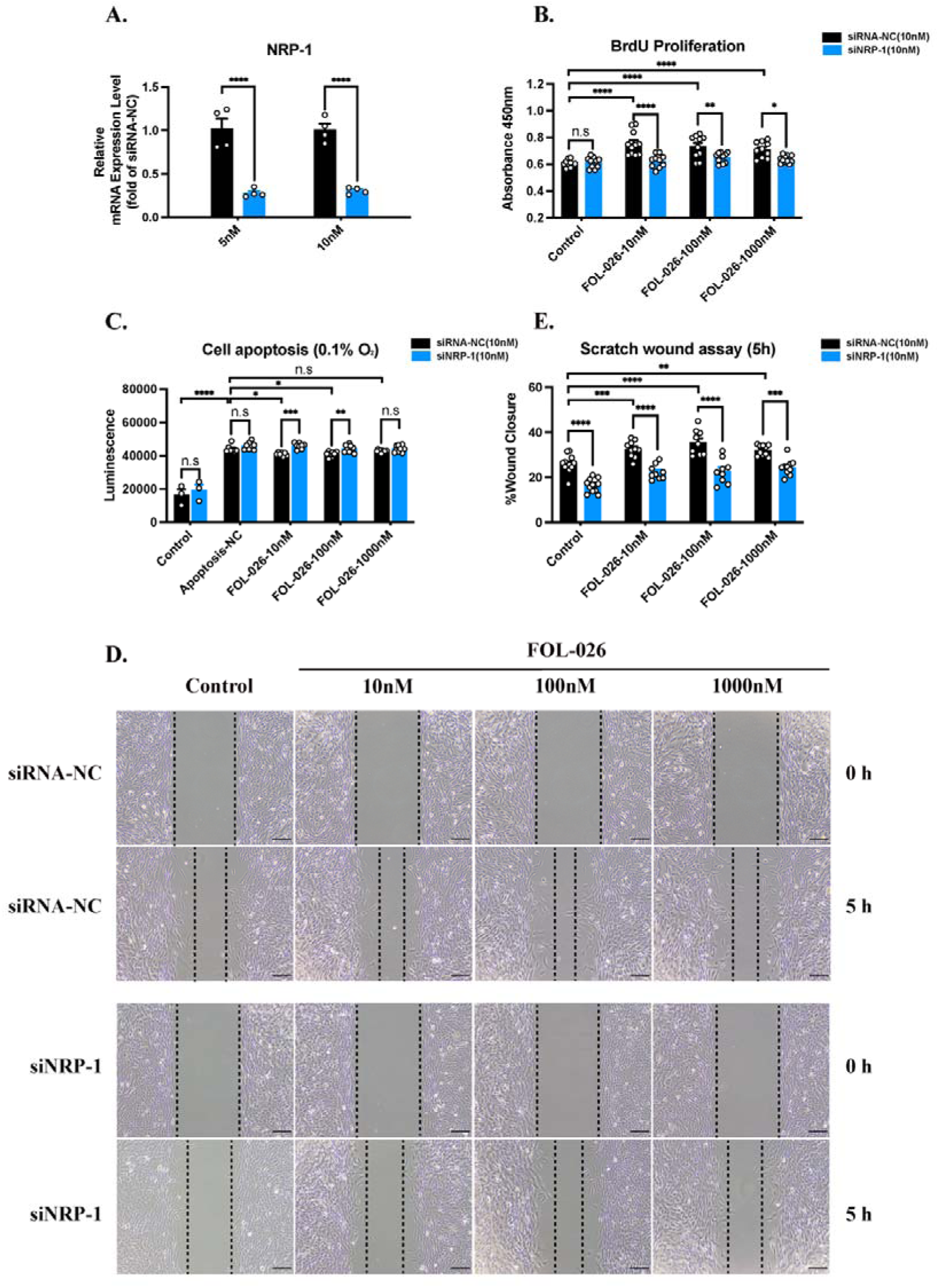
Effect of NRP-1 knock-down on arterial smooth muscle cells stimulated with FOL-026. **(A)** qRT-PCR analysis of NRP-1 mRNA expression in HCASMCs with small interfering RNA transfection for 48 hours (n=4). The assessment of BrdU incorporation (**B**, n=10-11) and caspase-3/-7 activation (**C**, n=3-10) in siRNA-NC/ siNRP-1-transfected HCASMCs treated with indicated concentration of FOL-026. Representative images of smooth muscle cells at 0 and 5 hours after induction of the wound are shown **(D)**. The wound closure rate **(E)** of HCASMCs in the siRNA-NC and siNRP-1 groups stimulated with FOL-026 peptide was quantified by image J software (n=9-12 per group). Scale bar =150μm in D. Data are presented as means±SEM and acquired from 3 to 4 independent replicate tests. Two-way ANOVA followed with Sidak’s multiple comparisons test for A, two-way ANOVA followed with Sidak’s and Dunnett’s multiple comparisons test for B, C and E.*P<0.05, **P<0.01, ***P<0.001, and ****P<0.0001.

### 3.7 Identification of genes and pathways regulated by FOL-026 in endothelial cells by RNA-seq

To further explore the pathways underlying the effects of FOL-026 on endothelial cells we performed RNA-seq analysis of HUVECs stimulated with increasing concentrations of FOL-026 for 48 hours (Fig.8A). Compared to control cells, there were 104 upregulated and 157 downregulated DEGs in HUVECs treated with 10nM of FOL-026, 51 upregulated and 104 downregulated DEGs in HUVECs treated with 100nM of FOL-026, and 112 upregulated and 115 downregulated DEG in HUVECs treated with 1000nM (Supplementary Fig.S6A). Among them, 28 DEG genes were overlapped between all concentration of FOL-026 compared to the controls. In line with the experimental studies, GO analysis revealed that the DEGs mostly associated with biological processes involved in tissue injury and repair including extracellular matrix organization, positive regulation of angiogenesis, wound healing and intrinsic apoptotic signaling pathway (Fig.8B). Furthermore, KEGG analysis showed that DEGs induced by FOL-026 were involved in protein digestion and absorption, extracellular matrix-receptor interactions, as well as the PI3K-Akt signaling pathway (Fig.8C). To visualize dynamic gene changes in response to increasing concentrations of FOL-026, DEGs were grouped into 12 clusters using Mfuzz package (Fig.8D). We used cluster 1, 4 and 5, which displayed marked increase or reduction in gene expression in response to FOL-026, to construct PPI network. As there was an attenuation of the effects of FOL-026 at the highest concentration in some of cell culture experiments we also included cluster 9, that was characterized by a decrease in gene expressions at 10nM and 100nM of FOL-026 followed by an increase at 1000nM concentration, in the PPI analysis. Based on the DEGs of different clusters, a PPI network with 167 nodes and 279 edges was established (Fig.8E). Finally, the top 10 hub genes including BCL2 like 1 (BCL2L1), ATM serine/threonine kinase (ATM), mitogen-activated protein kinase 3 (MAPK3), RB transcriptional corepressor 1(RB1), BCL2 like 11(BCL2L11), centromere protein E (CENPE), kinesin light chain 1 (KLC1), kinesin family member 3A (KIF3A), kinesin family member 3B (KIF3B) as well as CCAAT enhancer binding protein alpha (CEBPA) were identified using maximal clique centrality (MCC) method in Cytoscape software with cytohubba plug-in (Fig.8F and Supplementary Fig.S6C).

**Figure 8.**
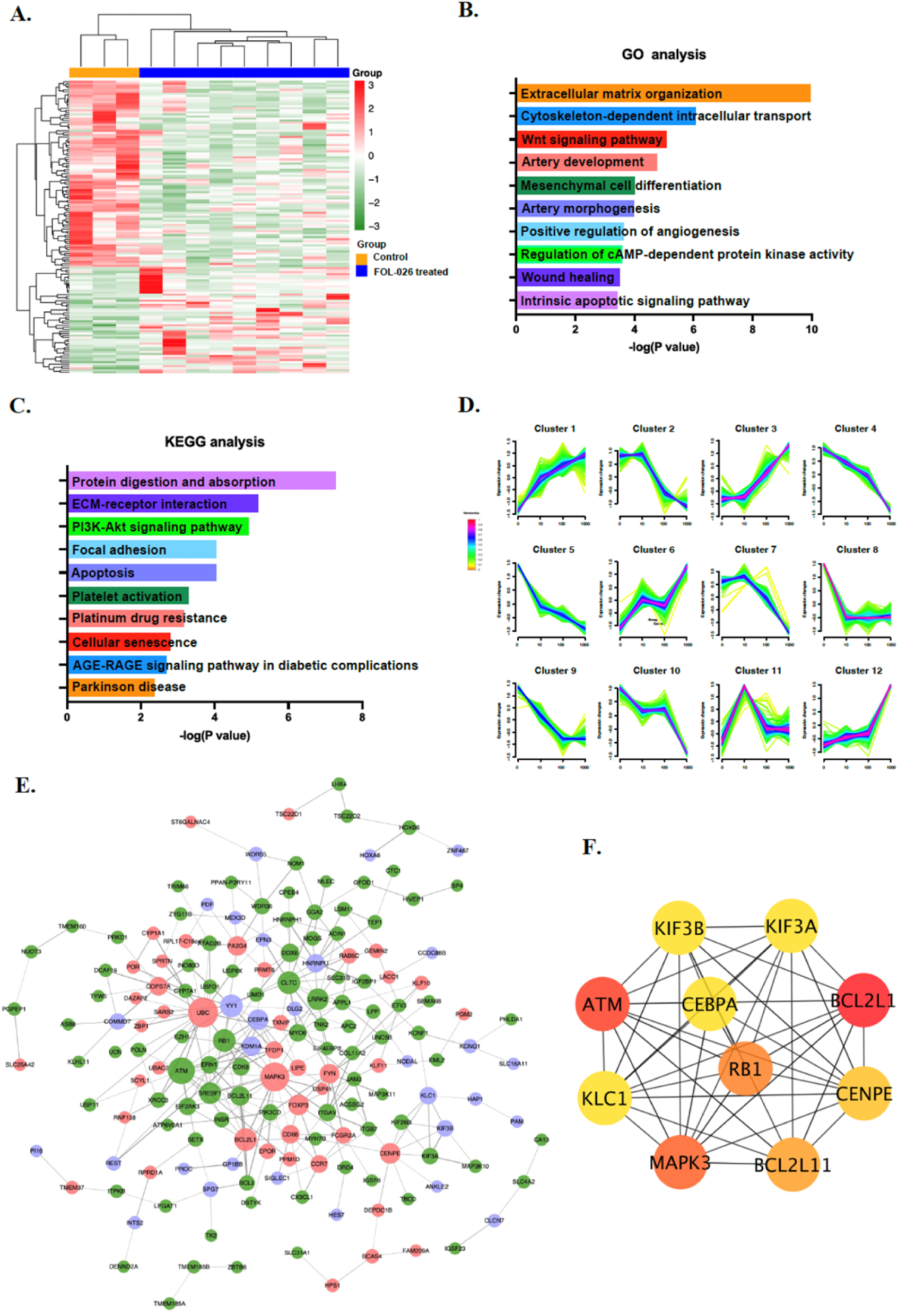
Key genes and pathways regulated by FOL-026 in endothelial cells. RNA-seq was performed on HUVECs treated with FOL-026 for 48 hours. The heatmap display the differently expression genes (DEGs) with *p* value <0.05 **(A)**. Gene ontology (GO) and Kyoto encyclopedia of genes and genomes (KEGG) enrichment analysis for DEGs. The top 10 biological process terms of GO analysis **(B)** and top 10 pathways of KEGG analysis are shown **(C)**. The DEGs were grouped into 12 clusters with Mfuzz package to describe the dynamic changes in response to increasing concentration of FOL-026 **(D)**. The DEGs (cluster 1, 4, 5 and 9) were used to construct protein - protein interaction (PPI) network **(E)** using Search Tool for the Retrieval of Interacting Genes (STRING) database. Red and green dots represent DEGs that were upregulated (cluster 1) and downregulated (cluster 4 and 5) with increasing concentration of FOL-026, respectively. Purple dots represent DEGs that were downregulated gradually with FOL-026 concentrations ranging from 0-100nM, while DEGs were upregulated at 1000nM compared to100nM (cluster 9). The top 10 hub genes were screened by maximal clique centrality (MCC) using cytohubba plug-in **(F)**. The color ranging from red (most crucial) to yellow (crucial) was used to describe the significance of hub genes.

### 3.8 Identification of candidate genes and pathways regulated by FOL-026 in arterial smooth muscle cells by RNA-seq

RNA-seq analysis of HCASMCs treated with different concentrations of FOL-026 also revealed distinct differences in gene expression between FOL-026-treated cells and controls (Supplemental Fig.7A). There were 62, 71 and 80 upregulated DEGs and 109, 132 and 184 downregulated DEGs in HCASMCs incubated with 10nM, 100nM and 1000nM of FOL-026, respectively compared with control cells (Supplementary Fig.S8A). Out of these, 36 genes were differentially expressed at all concentrations of FOL-026 as compared to the controls (Supplementary Fig.S8B). GO analysis revealed that the DEGs were mainly related with fatty acid and phospholipid metabolic process, the regulation of cell adhesion, wound healing, and angiogenesis (Supplemental Fig.7B). The activation of PI3K-Akt signaling by FOL-026 found by western blot assay was confirmed by KEGG analysis (Supplemental Fig.7C). Next, we classified all DEGs into 12 clusters with different modes of gene expression changes (Supplemental Fig.7D). The DEGs (cluster 1, 2, 4, 5, 9 and 12) were used to construct the PPI network for exploring interactions between pivotal DEGs by FOL-026 stimulation. To identify patterns of gene expression more clearly in the PPI network containing 475 nodes and 1330 edges were separated into two PPIs (Supplemental Fig. 7E and 7F). The PPI network based on DEGs from cluster 1, 2 and 9, whose expressions were consistently increased or decreased in response to FOL-026, is displayed in supplemental Fig.7E. The PPI network based on cluster 4, 5 and 12, where a dose-dependent increase or decline in gene expression at the two lower concentrations of FOL-026 were followed by an altered trend at the 1000nM concentration, is shown in supplemental Fig.7F. For hub gene identification, the nodes were calculated with MCC scoring method with the top 10 hub genes included kinase insert domain receptor (KDR, also known as VEGFR-2), CD34 molecule (CD34), KIT proto-oncogene, receptor tyrosine kinase (KIT), PECAM1, von Willebrand factor (VWF), C-X-C motif chemokine receptor 4 (CXCR4), platelet derived growth factor subunit B (PDGFB), fms related receptor tyrosine kinase 4 (FLT4, also known as VEGFR-3), placental growth factor (PlGF) and bone morphogenetic protein 4 (BMP4) (Supplemental Fig.7G and Supplementary Fig.S8C).

## 4. Discussion

We demonstrate here that the osteopontin-derived FOL-026 peptide stimulates migration and proliferation of HUVECs and HCASMCs. Furthermore, it also protects against apoptosis induced by hypoxia, oxLDL, and activation of the Fas receptor in these cells and stimulates the *in vitro* formation of endothelial tubes. These observations provide evidence for a pro-angiogenic effect of FOL-026. Earlier experimental and clinical studies have shown that treatment with the FOL-026 analogue FOL-005 as well as osteopontin^4,18^ stimulates hair growth but the mechanisms responsible for this effect have been unclear. Yano and coworkers^9^ were first to demonstrate that the transition of resting hair follicles into the anagen growth phase is preceded by an increased perifollicular vascularization and that this is activated by an increased expression of VEGF in follicular keratinocytes in the outer root sheath of the hair follicle and FOL-005 has been shown to also accumulate around and affect these cells.^19^ Treatment with VEGF blocking antibodies inhibited both perifollicular vascularization and hair growth. The critical role of perifollicular vascularization in stimulation of hair growth has subsequently been confirmed by several other investigators who showed that other growth factors belonging to the VEGF family, such as PlGF and HGF, have similar roles in the regulation of hair growth.^10–13,20^ Minoxidil, a drug approved for treatment of hair loss, is also known to function by stimulating perifollicular vascularization.^21^ Collectively, these observations argue that FOL-005 stimulates hair growth through stimulation of perifollicular vascularization. Since FOL-026 also activated the expression of VEGF in HCASMCs it is a possibility that the action of the peptide is indirect and depends on enhancing VEGF expression in dermal cells.

The receptors that mediate the effect of the VEGF growth factor family (VEGF receptor 1 and 2 for VEGFA and PlGF, and c-Met for HGF) use NRP-1 as co-receptor.^22^ These growth factors bind simultaneously to NRP-1 and the respective growth factor receptor resulting in formation of receptor complexes activating intracellular signals. In this context it is interesting to note that FOL-26 not only has similar biological effects as the members of the VEGF growth factor family, but that it also binds to the same site in NRP-1 as VEGFA. The mechanisms through which NRP-1 contributes to VEGF signaling are complex and remains to be fully characterized, but are known to involve dimerization with other receptors.^23^ Thus, while NRP-1 lacks an intracellular kinase domain of its own it can transmit signals by recruiting and dimerizing other receptor kinases. In our study, silencing NRP-1 expression by siRNA blunted the effect of FOL-026 on HUVEC and HCASMC providing evidence for a functional role of NRP-1 in mediating the effects of FOL-026. However, further studies are required to elucidate the exact mechanisms involved in NRP-1-mediated cell activation by FOL-026 including the attenuation of effects observed at higher concentration in some of the experiments. One possibility could be that FOL-26 activates dimerization of NRP-1 with other receptor kinases in the same way as VEGF family growth factors do. In this context it is interesting to note that FOL-026, like the VEGF growth factors, activates ERK1/2 and AKT. As discussed above it is also possible that the primary effect of FOL-026 is to activate the expression of VEGF in cells and that this growth factor in turn is responsible for the effects on HUVECs and HCASMCs proliferation, migration, and viability.

Members of the VEGF growth factor family play critical roles in the formation and maintenance of blood vessels both during development and in adult tissues.^24^ Most focus has been on development of anti-VEGF therapies for treatment of cancer but recently their importance in vascular repair and protection against cardiovascular disease has also received attention. In a population cohort of 4,742 individuals with 20-years follow up we found that a low ability to respond with expression of PlGF in response to metabolic stress was associated with an increased risk of myocardial infarction, stroke, and cardiovascular death.^17^ Analyzing atherosclerotic plaques removed at carotid surgery we also found that plaques with a high content of PlGF had a more stable phenotype and subjects with such plaques had a lower risk of future cardiovascular events.^17^ Moreover, using a Mendelian Randomization approach in more than 30,000 subjects Folkersen and coworkers found evidence for a protective role of PlGF in coronary heart disease.^25^ Lack of sufficient vascular repair capacity remains a relatively unexplored risk factor for cardiovascular disease, but it is conceivable that such a condition may increase the risk of thrombosis due to poor endothelial repair. This may be of particular importance for endothelial erosions, a phenomenon that is believed to be the cause of as much as one third of all myocardial infarctions.^26,27^ However, insufficient repair capacity may also contribute to an increased risk of plaque rupture due to a decreased ability to maintain the strength of the plaque fibrous cap. There is evidence that this phenomena may be of particular importance in subjects with diabetes.^28^ Although the risk of myocardial infarction and stroke has markedly decreased in subjects with diabetes during recent years it has only done so to the same extent as in the rest of the population leaving affected subjects at a 2-fold higher risk.^29^ This suggests that the pathophysiological mechanisms through which diabetes causes cardiovascular disease remains to be targeted by medical interventions.^30,31^ Biomarkers reflecting cellular stress are elevated in diabetes.^32^ Subjects with diabetes also have an impaired capacity of endothelial repair through bone marrow-derived endothelial progenitor cells as well as reduced levels of growth factors required for the activation of these cells.^33–35^ Moreover, atherosclerotic plaques in subjects with diabetes have thinner fibrous caps and contain less amounts of fibrous tissue.^28,36^ The possibility of developing treatments that enhance vascular repair is therefore of potential interest. Both our cell culture studies and RNA-seq analysis show that FOL-026 activates processes involved in vascular repair indicating that FOL-026 t represents an interesting potential therapeutic candidate in this context. Therapies that stimulate angiogenesis and vascular repair may also have beneficial effects in conditions such as diabetic foot ulcers, patency of vascular grafts, peripheral artery disease, and tissue reconstructions.

The C-terminal end of FOL-005 and FOL-026 contains the amino acid sequence SVVYGLR. The possible biological role of this peptide sequence has attracted attention since it becomes exposed following thrombin cleavage of osteopontin.^2,37^ Synthetic SVVYGLR peptide has been shown to stimulate proliferation and migration of endothelial cells^38^ as well as to stimulate the functional regeneration of oral skeletal muscle.^39^ The effect of the SVVYGLR peptide has been shown to be mediated through binding to α4 and α9 integrins.^40,41^ In the present study we found that the C-terminal amino acids leucine (L) and arginine (R) are critical for binding of FOL-026 to NRP-1 and that downregulation of NRP-1 blocks the stimulatory effects of FOL-026 on endothelial cells. Using the Cell Microarray technology (Retrogenix) we were unable to identify binding of FOL-026 to α4 and α9 integrins (data not shown). It is clear that at least part of the SVVYGLR amino acid sequence plays an important role in mediating the biological effects of FOL-026 but this will require further investigation since different receptors appear to be involved.

There are some limitations of the present study that should be considered. The exact mechanisms through which NRP-1 mediates the effects of FOL-026 need to be fully elucidated as well as the reason for the bell-shaped dose-response curve observed in some experiments. Since NRP-1 is a coreceptor for the VEGF family of growth factors and the biological effects of FOL-026 are very similar to that of these growth factors it is likely that FOL-026 in some way interacts with the VEGF and/or c-Met receptors. Moreover, further studies in appropriate animal models and clinical studies are required to confirm the angiogenic properties of FOL-026 as well as the importance of these for the effect of FOL-005 on hair growth.

## Conclusions

The present study shows that the osteopontin-derived peptide FOL-026 binds to same binding site in NRP-1 as VEGF and the biological effects of FOL-026 are like those of the VEGF family of growth factors. They also suggest that these peptides may have therapeutical potential in conditions that require vascular repair or enhanced angiogenesis including vulnerable atherosclerotic lesions, salvage of ischemic tissues and healing of chronic ulcers.

## Acknowledgements

We are grateful to Dualsystems Biotech AG, Schlieren Switzerland for using LRC-TriCEPS technology to identify FOL-005 receptor candidates.

## Nonstandard Abbreviations and Acronyms

NRP-1: neuropilin-1
VEGF: vascular endothelial growth factor
HGF: hepatocyte growth factor
HUVEC: human umbilical vein endothelial cells
HCASMC: human coronary artery smooth muscle cells
siRNA: small interfering RNA
ROS: reactive oxygen species
oxLDL: oxidized low-density lipoprotein
sFasL: soluble Fas ligand
*RNA*-seq: *RNA* sequencing
DEGs: differentially expressed genes
GO: gene ontology
KEGG: kyoto encyclopedia of genes and genomes
BP: biological processes
PPI: protein-protein interaction
STRING: search tool for the retrieval of interacting genes
MST: microscale thermophoresis
SEM: standard error of mean
VEGFR-2: vascular endothelial growth factor receptor-2
c-Met: MET proto-oncogene, receptor tyrosine kinase/hepatocyte growth factor receptor
TNFα: tumor necrosis factor α
IL-6: interleukin 6
MMP2: matrix metalloproteinase-2
MMP3: matrix metalloproteinase-3
COL1A1: collagen type I α 1 chain
COL1A2: collagen type I α 2 chain
MCC: maximal clique centrality
BCL2L1: BCL2 like 1
ATM: ATM serine/threonine kinase
MAPK3: mitogen-activated protein kinase 3
RB1: RB transcriptional corepressor 1
BCL2L11: BCL2 like 11
CENPE: centromere protein E
KLC1: kinesin light chain 1
KIF3A: kinesin family member 3A
KIF3B: kinesin family member 3B
CEBPA: CCAAT enhancer binding protein alpha
CD34: CD34 molecule
KIT: KIT proto-oncogene, receptor tyrosine kinase
PECAM1: platelet endothelial cell adhesion molecule-1
VWF: von Willebrand factor
CXCR4: C-X-C motif chemokine receptor 4
PDGFB: platelet derived growth factor subunit B
FLT4: fms related receptor tyrosine kinase 4, as known as VEGFR-3
PlGF: placental growth factor
BMP4: bone morphogenetic protein 4

